# Affective enhancement of episodic memory is associated with widespread patterns of intrinsic functional connectivity across the adult lifespan

**DOI:** 10.1101/2022.03.31.485630

**Authors:** Yuta Katsumi, Matthew Moore

**Author notes:** **Correspondence:** Yuta Katsumi, PhD & Matthew Moore, PhD, /.

## Abstract

Subjectively arousing experiences tend to be better remembered than neutral ones. While numerous task-related neuroimaging studies have revealed the neural mechanisms associated with this phenomenon, it remains unclear how variability in the extent to which individuals show superior memory for subjectively arousing stimuli is associated with the *intrinsic* functional organization of their brains. Here, we addressed this issue using functional magnetic resonance imaging data collected at rest from a sample drawn from the Cambridge Centre for Ageing Neuroscience cohort (*N* = 269, 18-86 years). Specifically, we performed multi-voxel pattern analysis of intrinsic functional connectivity, an unbiased, data-driven approach to examine whole-brain voxel-wise connectivity patterns. This technique allowed us to reveal the most important features from the high-dimensional, whole-brain connectivity structure without a priori hypotheses about the topography and direction of functional connectivity differences. Behaviorally, both item and associative memory accuracy were enhanced for trials with affectively arousing (positive or negative) stimuli than those with neutral ones. Whole-brain multi-voxel pattern analysis of functional connectivity revealed that the affective enhancement of memory was associated with intrinsic connectivity patterns of spatially distributed brain regions belonging to several functional networks in the cerebral cortex. Post hoc seed-based brain-behavior regression analysis and principal component analysis of the resulting correlation maps showed that these connectivity patterns were in turn primarily characterized by the involvement of heteromodal association (dorsal attention, salience, and default mode) networks as well as select subcortical structures (striatum, thalamus, and cerebellum). Collectively, these findings suggest that the affective enhancement of episodic memory may be characterized as a whole-brain phenomenon, possibly supported by intrinsic functional interactions across several networks and structures in the brain.

## Introduction

Whether it is a wedding, college graduation, birth of a child, or death of a loved one, subjectively arousing experiences tend to be better remembered than neutral ones. Such affective enhancement of episodic memory has been well supported by scientific studies, with a wealth of empirical evidence identifying the neural mechanisms underlying this phenomenon (reviewed in Dolcos, Katsumi, Weymar, et al., 2017; Dolcos et al., 2020; Kensinger & Ford, 2020; LaBar & Cabeza, 2006). However, most of the prior studies in this domain have focused on relating brain activity and behavioral responses observed during the same task. Therefore, it remains unclear how variability in the extent to which individuals show superior memory for subjectively arousing stimuli is associated with the *intrinsic* functional organization of their brains. Clarification of this issue is important given that better understanding of the neural basis of enhanced memory for affectively arousing stimuli has translational implications. For instance, excessive (often involuntary) rumination of negative and distressing memories is known to be a hallmark of psychiatric disorders including major depression (Nolen-Hoeksema et al., 2008) and post-traumatic stress disorder (Berntsen & Rubin, 2014). Intrinsic neural signatures of the affective enhancement of memory, if they exist and can be reliably measured, may have utility as a biomarker to be targeted by preventive or therapeutic interventions designed to ameliorate the symptoms of affective disturbances.

Much of current knowledge about the neural basis of the affective enhancement of memory is based on studies utilizing various neuroimaging techniques, such as event-related functional magnetic resonance imaging (fMRI). In typical neuroimaging experiments of affective episodic memory, participants perform an encoding task in which they are presented with a series of stimuli varying in affective arousal (and valence) while their brain activity is recorded. After a delay, participants perform a retrieval task where they are presented with another set of stimuli—this time with some previously unseen information included—and are asked to correctly discriminate the old stimuli from the new ones. Participants are most commonly asked to recognize salient features of these stimuli (e.g., objects; a measure of *item* memory), while in some cases they are also asked to identify the contextual details associated with them (e.g., type of background, location on the screen, order of presentation; measures of *associative* memory). Comparisons between brain activity associated with subsequently remembered versus forgotten stimuli during encoding allow investigators to isolate and identify activation patterns related to successful encoding of stimuli (Paller & Wagner, 2002). By extension, comparing brain activity linked to successful encoding of more versus less arousing stimuli allows estimation of the effect of affective arousal on successful memory encoding activity (Dolcos, Katsumi, Denkova, et al., 2017). A similar approach can be used to examine brain activity measured at the time of retrieval.

Utilizing these experimental designs with healthy young adult participants, previous functional neuroimaging studies showed that successful affective episodic memory encoding and retrieval are associated with greater activation of an extensive set of brain regions, including those within the medial temporal, visual association, lateral prefrontal, lateral parietal, lateral temporal, and insular cortices (for meta-analyses, see Dahlgren et al., 2020; Murty et al., 2010). These regions are consistent with those identified by other, more general meta-analyses on affective processes, showing that processing of affective arousal involves a distributed network of brain regions from early sensory areas to heteromodal association areas of the cerebral cortex as well as subcortical structures (Hayes & Northoff, 2011; Lindquist et al., 2016; Satpute et al., 2015, 2019). Collectively, these findings suggest that the affective enhancement of episodic memory is likely not supported by computations localized to a specific region or a small set of regions, but is rather more accurately characterized as a whole-brain phenomenon.

To our knowledge, no published studies to date have investigated the relationship between intrinsic functional properties of the brain and individual differences in the affective enhancement of episodic memory. However, related evidence suggests that intrinsic functional connectivity may support affective memory biases exhibited behaviorally. For instance, functional connectivity at rest between the amygdala and the medial prefrontal cortex predicted the extent to which older adults preferentially remembered stimuli exhibiting facial movements prototypically associated with happiness versus anger in a subsequent task (Sakaki et al., 2013). In addition, intrinsic functional connectivity between the amygdala and the hippocampus was related to individual variation in the extent of memory enhancement by a task-irrelevant induction of stress (de Voogd et al., 2017). Because the functional architecture of the brain appears to be remarkably stable across task states (e.g., Cole et al., 2014; Finn et al., 2015; Gratton et al., 2016, 2018; Krienen et al., 2014; but see e.g., Geerligs et al., 2015), it is possible that the affective enhancement of episodic memory is related to functional connectivity patterns of widespread regions in the brain while it is not probed by external tasks (i.e., at rest).

The goal of the present study was to investigate the relationship between intrinsic functional connectivity and the affective enhancement of episodic memory across healthy adults. We sought to identify and characterize differences in intrinsic brain connectivity as a function of the affective enhancement of episodic memory using functional connectivity multi-voxel pattern analysis (MVPA) (Katsumi, Kondo, et al., 2021; Nieto-Castanon, 2020), an unbiased, data-driven approach to examine whole-brain intrinsic functional connectivity derived from fMRI data collected at rest. This technique allowed us to reveal the most important features from the high-dimensional, whole-brain connectivity structure without a priori hypotheses about the topography and direction of functional connectivity differences. We then compared the topography of our results with established canonical functional networks of the cerebral cortex (Yeo et al., 2011). This approach allowed us to characterize the patterns of significant functional connectivity differences in terms of a commonly used parcellation scheme to describe the functional organization of the cerebral cortex.

## Materials and Methods

### Participants

In this study, we analyzed behavioral and brain imaging data acquired from the Cambridge Center for Cognitive and Aging (Cam-CAN) sample, a population-representative cohort of cognitively normal adult participants aged 18 to 88 years (Shafto et al., 2014). Ethical approval for the Cam-CAN study was obtained from the Cambridgeshire 2 (now East of England-Cambridge Central) Research Ethics Committee and all participants gave written informed consent. We identified a subset of 312 participants who had complete affective episodic memory task data, structural MRI data, and fMRI data collected at wakeful rest (Taylor et al., 2017). From this initial pool of participants, we excluded 43 participants following preprocessing of fMRI data, resulting in the final sample of 269 participants (*M*_age_ = 52.03 ± 18.13, 130 males/139 females) for analyses; see below for exclusion criteria.

### Affective episodic memory task

Details of the affective episodic memory task are discussed elsewhere (Henson et al., 2016) and thus are only briefly reported here. Stimuli consisted of 160 images of everyday, affectively neutral objects overlaid on a yellow, square background (Smith et al., 2004), along with 120 images drawn from the International Affective Picture System (IAPS) (Lang et al., 1997). The IAPS images were classified as either positive, negative, or neutral (40 in each category), based on ratings of affective properties given by an independent sample of young (n = 12) and older (n = 7) adult participants (Henson et al., 2016). The mean valence and arousal ratings were significantly different in every pairwise comparison of the valence categories, with negative images being perceived as more arousing than positive ones.

The task was comprised of a Study phase and a Test phase (**Supplementary Figure S1**). The Study phase consisted of 120 trials split into 2 runs; each run lasted 10 minutes to complete. Each trial began with a neutral background scene presented for 2 seconds, after which an object appeared for 7.5 seconds either left or right and slightly above center. Participants were asked to respond with a key press when they had made a story in their mind that linked the object to the scene, and continue to elaborate that story until the scene and object disappeared. The intertrial interval was 0.5 seconds where a blank screen was presented. Participants were aware that some scenes would be pleasant or unpleasant but had not been told that their memory for the images would be tested later.

Participants proceeded to the Test phase of the task following a break of approximately 10 minutes. This phase consisted of 160 trials (120 “old” objects presented during the Study phase and additional 40 new objects), split into 4 runs; each run lasted approximately 20 minutes. Each trial began with a measure of *priming*; the masked, degraded version of an object appeared at the center of the screen and participants were asked to identify the object with a key press and to simultaneously name it aloud (or say “don’t know”). The pixel noise was then removed, and participants were asked to indicate whether they had seen the object during the Study phase and how confident they were in their decision, with possible choices being “sure new”, “think new”, “think studied”, “sure studied” (a measure of item memory). If participants indicated “think studied” or “sure studied”, then they were asked to say aloud whether the object had been paired with a positive, neutral, or negative background during the Study phase (or “don’t know” if they were unable to guess) (a measure of associative memory). Finally, participants were asked to verbally describe the recalled scene (the associated qualitative data are not reported here).

### Behavioral data analysis

By design, the affective memory task allows analyses of three different types of memory (priming, item memory, and associative memory) as well as the effect of affective content. Here, we focused on item and associative memory because the original study by the Cam-CAN group identified no significant effects of affective content on priming (Henson et al., 2016). Following these authors and also our prior work (Katsumi & Dolcos, 2018), we quantified memory accuracy with the *d’* measure of discriminability defined as the difference in inverse normal transformed probabilities of Hits and False Alarms, where *d’* = 0 indicates chance performance. Extreme values of 0 or 1 for *p*(Hits) and *p*(False Alarms) were adjusted using an empirical log-linear approach. Hits and False Alarms were collapsed across “sure” and “think” levels of confidence for item memory due to the small number of low confidence judgments. False Alarms for associative memory were defined as incorrectly endorsing a given (e.g., negative) background condition when the correct answer should have been one of the other two (e.g., positive or neutral). Memory accuracy was analyzed at the group level via a one-way repeated-measures analysis of variance (ANOVA) with Affective Content (negative, positive, neutral) as a factor, separately for each memory type. Post hoc comparisons were performed with Bonferroni corrected *p* values (*p*_bonf_) to examine the direction of effects.

To quantify the magnitude of the affective enhancement of memory in each participant, we computed the difference in *d’* across conditions as follows, separately for item and associative memory:

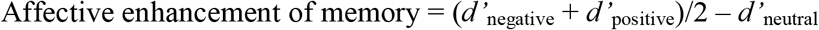

That is, more positive values indicate better memory for images with higher subjectively arousing content, regardless of valence, compared with neutral images.

### MRI data acquisition and preprocessing

Structural and functional MRI data were acquired on a 3T Siemens Tim Trio system using a 32-channel head coil. A high-resolution 3D T1-weighted anatomical image was acquired using a MPRAGE sequence: Repetition time (TR) = 2250 ms, echo time (TE) = 2.99 ms, inversion time (TI) = 900 ms, flip angle = 9°, field of view (FOV) = 256 mm × 240 mm × 192 mm, 1 mm isotropic voxels, GRAPPA acceleration factor = 2, acquisition time = 4 minutes 32 seconds. A functional echo planar imaging scan was acquired while participants rested with eyes closed for 8 minutes 40 seconds: TR = 1970 ms, TE = 30 ms, flip angle = 78°, FOV = 192 mm × 192 mm, 32 axial slices, voxel size = 3 mm × 3 mm × 4.44 mm.

Preprocessing of MRI data was performed in SPM12 (v7487; Wellcome Department of Cognitive Neurology, London, UK). The first four functional images were discarded to allow scanner equilibrium effects. Functional images were then spatially realigned and resliced relative to the mean volume to correct for between-scan motion (using a six-parameter rigid body transformation) and were also slice-time corrected relative to the first slice. Finally, these images were resampled to 2 mm isotropic voxels during direct normalization to the MNI152 space, after which they were spatially smoothed using a 6 mm Gaussian kernel, full-width at half-maximum. To address the spurious correlations induced by head motion, outlier scans were identified using the Artifact Detection Tools (ART, www.nitrc.org/projects/artifact_detect/). Specifically, a scan was defined as an outlier/artifact scan if (1) the global signal intensity was > 3 SD above the mean signal or (2) composite head movement exceeded 0.5 mm from the previous scan (Katsumi, Kondo, et al., 2021; Power et al., 2014). The composite head movement was computed by first converting six rotation/translation head motion parameters into another set of six parameters characterizing the trajectories of six points located on the center of each of the faces of a bounding box around the brain. The maximum scan-to-scan movement of any of these points was then computed as the single composite movement measure (Chai et al., 2016). Each outlier scan was represented by a single regressor in the general linear model for each participant, with a value of 1 for each outlier scan and 0s elsewhere. Participants with less than 5 min worth of fMRI data after discarding the outlier scans were excluded from all analyses (n = 43). The mean of scan-to-scan motion across all the remaining participants was 0.21 ± 0.06 mm.

In addition, physiological and other spurious sources of noise were estimated and regressed out using the anatomical CompCor method (Behzadi et al., 2007), as implemented in the CONN toolbox v21a (Whitfield-Gabrieli & Nieto-Castanon, 2012). The normalized anatomical image for each participant was segmented into gray matter (GM), white matter (WM), and cerebrospinal fluid (CSF) masks in SPM12. To minimize partial voluming with GM, WM and CSF masks were eroded by one voxel, resulting in substantially smaller masks than the original segmentations (Chai et al., 2012). The eroded WM and CSF masks were then used as noise ROIs. Signals from the WM and CSF noise ROIs were extracted from the unsmoothed functional images to avoid additional risk of contaminating WM and CSF signals with GM signals. As part of the denoising procedure, the following nuisance variables were regressed out from the data: five principal components of the signals from WM and CSF noise ROIs, residual head motion parameters (three rotation and three translation parameters, along with their first-order temporal derivatives), each outlier/artifact scan, and linear trends. Finally, a temporal bandpass filter of 0.008-0.09 Hz was applied to the residual time series.

### Functional connectivity multi-voxel pattern analysis (MVPA)

**Figure 1** summarizes the analytical pipeline of this study. Using preprocessed time series data, we first identified brain regions whose patterns of whole-brain functional connectivity were associated with the affective enhancement of memory using functional connectivity MVPA (Nieto-Castanon, 2020). The utility of this method has been established in numerous prior studies examining brain-behavior relationships in healthy and clinical populations (e.g., Arnold Anteraper et al., 2019, 2020; Beaty et al., 2015; Guell et al., 2020; Katsumi, Kondo, et al., 2021; Morris et al., 2021; Muehlhan et al., 2020; Takamiya et al., 2021; Whitfield-Gabrieli et al., 2016). Features of interest were extracted at the first level using principal component analysis (PCA). In the first step, for each participant, 64 subject-specific PCA components were identified to characterize its pattern of correlations with all the other voxels in the brain. This step generated a low-dimensional representation of whole-brain correlation structure similar to other established data-driven approaches (e.g., Calhoun et al., 2012). In the second step, jointly across all participants but separately for each voxel, seven strongest components were retained from a PC decomposition of the between-subjects variability in voxel-to-voxel connectivity maps. This resulted in seven values per voxel that represented the entire connectivity pattern between a given voxel and the rest of the brain for each participant. The number of components (*N_c_* = 7) was determined by the participant-to-component ratio as well as the individual and cumulative explained variance for each *n* component (Morris et al., 2021) (see **Supplementary Materials and Methods**). We analyzed simultaneously these seven component maps by a single second-level *F*-test separately for the affective enhancement of item and associative memory to identify voxels whose functional connectivity patterns varied as a function of these behavioral measures. All MVPA results were thresholded using a voxel-wise intensity threshold of *p* < .001 with cluster-level correction for family-wise error rate (FWER) at *p* < .05, controlling for age, sex, and mean composite motion (calculated without outlier scans) as covariates.

**Figure 1.**
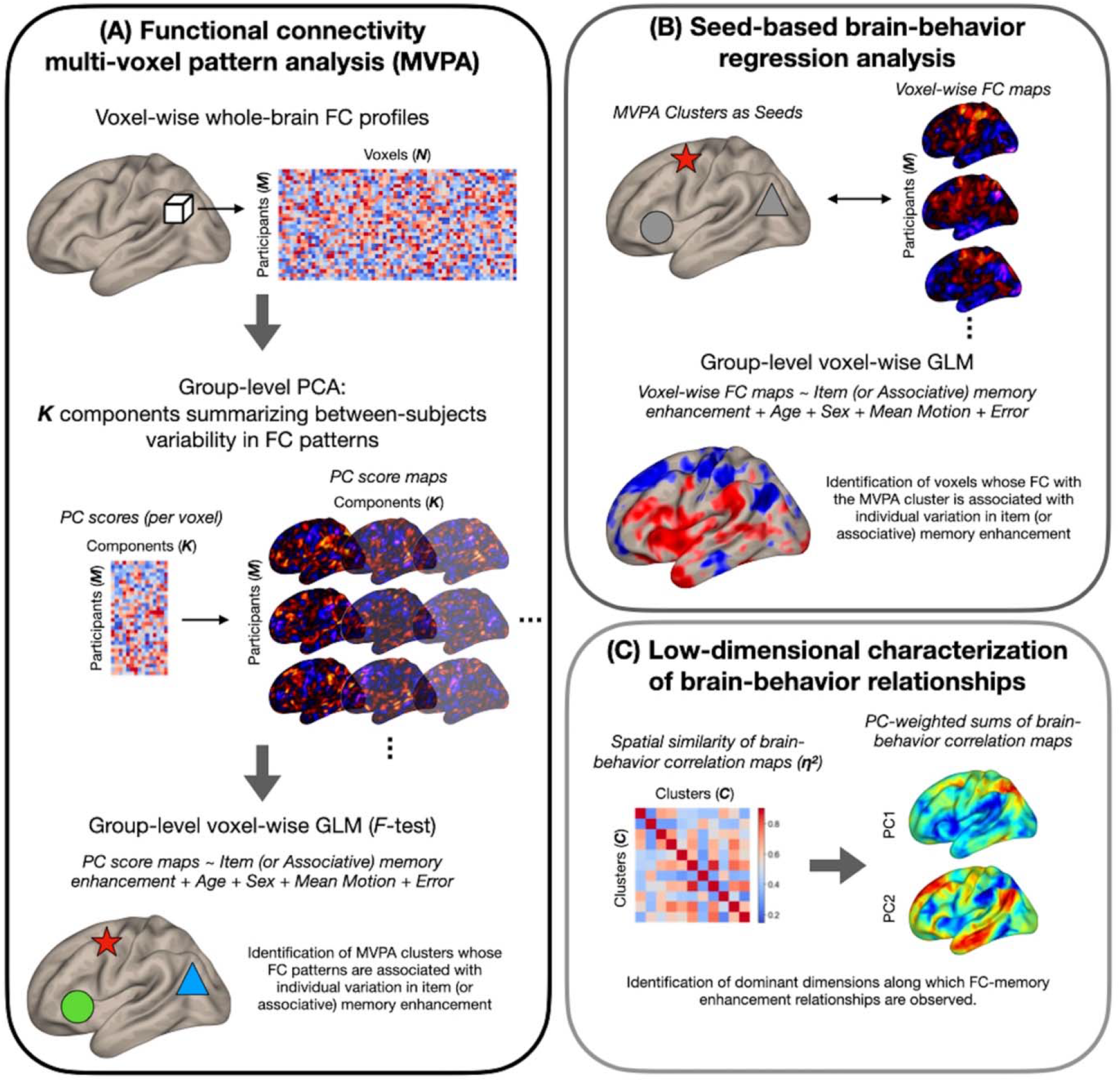
Schematic diagram of the analytical pipeline.

### Post hoc seed-based regression analysis

Upon identification of the significant MVPA clusters, we conducted post hoc seed-based regression analysis to characterize functional connectivity differences as a function of the affective enhancement of memory. For each participant, we calculated Pearson’s correlation coefficients between the mean time course of each significant MVPA cluster (calculated using unsmoothed data within each cluster) and that of all other voxels in the brain. Individual voxelwise connectivity maps were *z*-transformed and submitted to second-level multiple regression analyses to investigate the extent of correlation between functional connectivity and behavioral measures of memory enhancement. Results from this post hoc analysis were thresholded using a voxel-wise intensity threshold of *p* < .001 with cluster-level correction for FWER at *p* < .05, controlling for age, sex, and mean composite motion (calculated without outlier scans) as covariates.

### Low dimensional decomposition of brain-behavior correlations

Whole-brain MVPA and post hoc regression analysis revealed the involvement of widespread networks and regions in the brain (see **Results**). To facilitate the interpretation of these results, we performed low dimensional decomposition of second-level brain-behavior regression maps resulting from all 11 MVPA clusters. Specifically, we first quantified the extent of voxel-wise spatial similarity across all 11 maps by η^2^ (Cohen et al., 2008). This yielded a 11 × 11 affinity matrix quantifying pairwise similarity in the spatial topography of brain-behavior correlation, which was submitted to PCA. For each resulting PC, we computed the sum of all 11 brain-behavior regression maps weighted by the respective PC scores (Katsumi et al., 2022; Zhang et al., 2019). These component-weighted sum of brain-behavior regression maps revealed dominant axes along which the spatial topography of brain-behavior relationships can be summarized.

### Functional network-based characterization of second-level results

To facilitate the topographic characterization of functional connectivity and correlation differences, we compared the topography of relevant second-level statistical maps with the macroscale organization of intrinsic functional connectivity. Specifically, we calculated the extent of voxel-wise spatial overlap between these second-level statistical maps and large-scale functional networks as defined by a common parcellation of the cerebral cortex (Yeo et al., 2011). This procedure identified the relative proportion of voxels in each contributing map belonging to each of the seven canonical networks (i.e., visual, somatomotor, dorsal attention, salience, limbic, frontoparietal, and default mode). Here, we adhere to the original and conventional use of these network labels, although we acknowledge that the “default” and “limbic” networks are not always distinguished in the literature (e.g., Kong et al., 2019) and that both networks contain agranular, limbic tissue (Katsumi et al., 2022; Kleckner et al., 2017). In addition, anatomical labels of each suprathreshold cluster were examined and reported using the Harvard-Oxford cortical and subcortical atlas (Desikan et al., 2006; Frazier et al., 2005; Goldstein et al., 2007; Makris et al., 2006) as well as the AAL atlas (Tzourio-Mazoyer et al., 2002).

## Results

### Affective enhancement of item and associative memory performance

Item and associative memory accuracy data are reported elsewhere based on a superset of the Cam-CAN cohort (Henson et al., 2016). Here, we briefly report the results based on the sample of this study, which are consistent with Henson et al.’s (2016) findings. A repeated-measures ANOVA on item memory accuracy revealed a significant effect of Affective Content: *F*(2,536) = 15.66, *p* ≤ .001, η^2^_p_ = .055. Item memory was better for both positive (*M* = 2.76, *SD* = 0.73) and negative (*M* = 2.70, *SD* = 0.79) trials than neutral ones (*M* = 2.63, *SD* = 0.75): Positive vs. neutral *t* = 5.60, *p*_bonf_ ≤ .001, *d* = 0.34; negative vs. neutral *t* = 2.92, *p*_bonf_ ≤ .011, *d* = 0.18. Item memory was better for positive than for negative trials: *t* = 2.68, *p*_bonf_ ≤ .023, *d* = 0.16. Turning to associative memory accuracy, a repeated-measures ANOVA yielded an even larger effect of Affective Content: *F*(2,536) = 384.52, *p ≤* .001, η^2^_p_ = .056. Specifically, associative memory was again better for both positive (*M* = 1.25, *SD* = 0.62) and negative (*M* = 1.65, *SD* = 0.74) trials than neutral ones (*M* = 0.97, *SD* = 0.60): Positive vs. neutral *t* = 11.57, *p*_bonf_ ≤ .001, *d* = 0.71; negative vs. neutral *t* = 27.61, *p*_bonf_ ≤ .001, *d* = 1.68. Associative memory was better for negative than for positive trials: *t* = 16.05, *p*_bonf_ ≤ .001, *d* = 0.98.

### Intrinsic functional connectivity and its relation to item memory enhancement

Whole-brain functional connectivity MVPA revealed three clusters showing significant associations with the magnitude of the affective enhancement of item memory: Left frontal pole (peak MNI coordinates: −20, 68, −2; cluster size *k* = 42), right frontal pole (26, 54, 36; *k* = 44), and left postcentral gyrus (−38, −24, 38; *k* = 51) (**Figure 2**, upper panel). These clusters spatially overlapped the most with the default mode, frontoparietal, and somatomotor networks, respectively. To further characterize intrinsic functional connectivity differences as a function of item memory enhancement, we performed post hoc seed-based brain-behavior regression analysis using as seeds all three clusters identified by MVPA above. This analysis revealed widespread areas of the cerebral cortex whose functional connectivity with the MVPA seed regions showed positive or negative correlations with item memory enhancement (**Figure 2**, lower panels). Although overall these results involved regions from several functional networks in the cerebral cortex, the topography of suprathreshold voxels in brain-behavior regression maps revealed bilateral localization of effects for each MVPA cluster that were characteristic of specific networks. For instance, item memory enhancement was negatively associated with functional connectivity between the left frontal pole seed (default mode) and regions including bilateral dorsomedial prefrontal cortex, posterior cingulate cortex, angular gyrus, and lateral temporal cortex, thus largely representing the same network. Item memory enhancement was also associated with functional connectivity between the left postcentral gyrus (somatomotor) and bilateral medial prefrontal cortex (default mode; positive) as well as bilateral supramarginal gyrus (salience; negative). Similar effects were observed with functional connectivity between the right frontal pole (frontoparietal) and bilateral middle/inferior frontal gyrus, insula, and supramarginal gyrus (frontoparietal/salience; positive) as well as bilateral ventromedial prefrontal cortex, posterior cingulate cortex, and retrosplenial cortex (default mode; negative). Outside the boundaries of the cerebral cortical functional networks, item memory enhancement was associated with functional connectivity between the left frontal pole seed (default mode) and bilateral clusters in mid to posterior Crus I and Crus II in the cerebellar cortex (**Supplementary Figure S2**).

**Figure 2.**
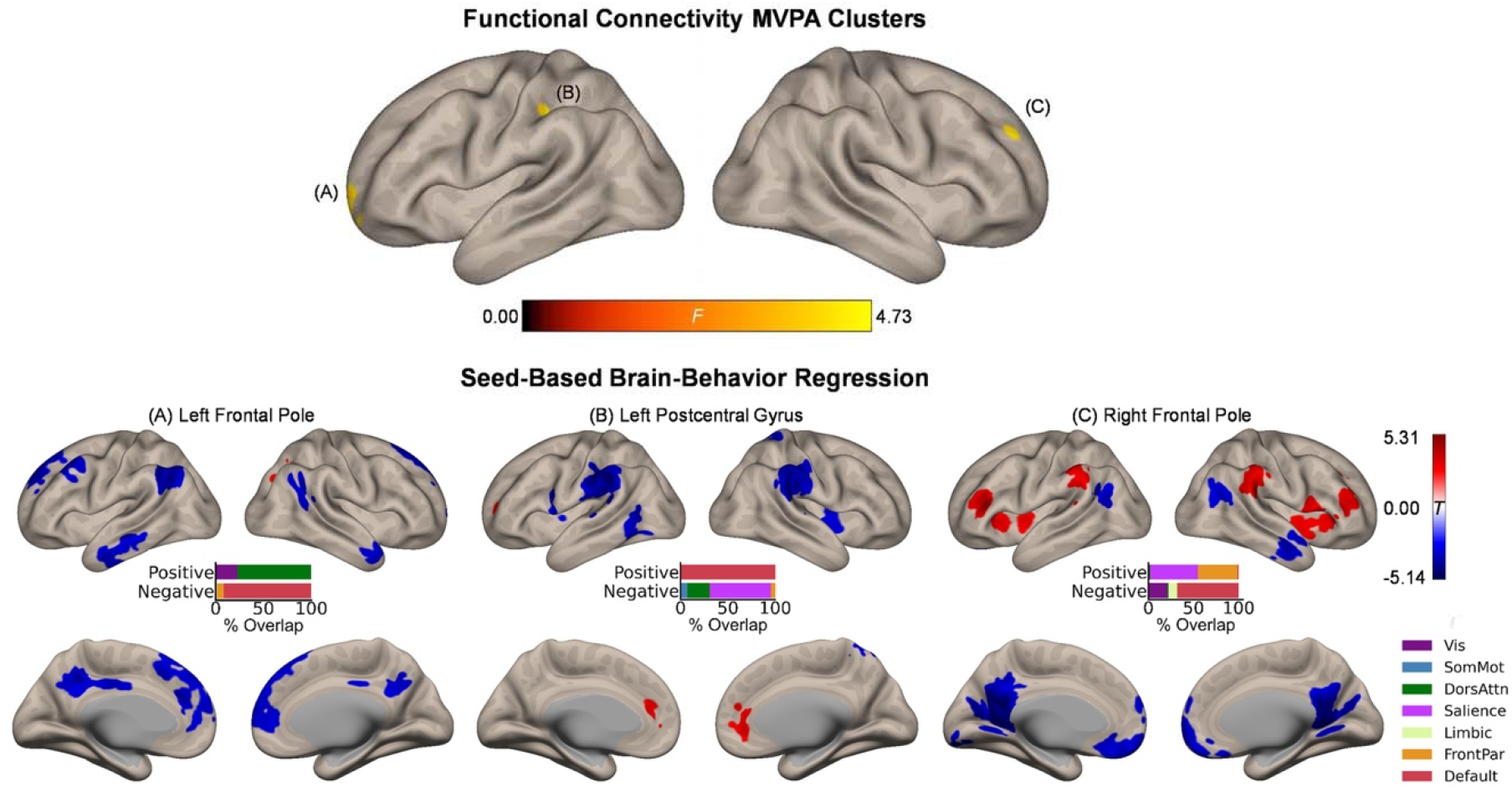
Intrinsic functional connectivity correlates of item memory enhancement. Upper panel: Clusters identified in the functional connectivity MVPA as being significantly associated with item memory enhancement. Lower panels: Whole-brain correlation maps between seed connectivity and item memory enhancement. Bar graphs show the percentage of overlap between the areas identified in the correlation maps and canonical functional networks of the cerebral cortex (Yeo et al., 2011). Note that the voxel-wise intensity threshold for the correlation maps corresponds to |*t*| = 3.12 (*p* ≤ .001).

### Intrinsic functional connectivity and its relation to associative memory enhancement

Whole-brain MVPA revealed eight clusters showing significant associations with the magnitude of the affective enhancement of associative memory, including right inferior frontal gyrus (44, 26, 14; *k* = 37), left precentral/middle frontal gyrus (−32, −6, 56; *k* = 74), right precentral gyrus (40, −14, 56; *k* = 82), precuneus (8, −54, 46; *k* = 64) left superior temporal gyrus (−56, −6, 2; *k* = 93), left (−42, −72, −6; *k* = 77) and right (48, −70, −8; *k* = 48) lateral occipital cortex, and lingual gyrus (−10, −76, 2; *k* = 63) (**Figure 3**, upper panel). Spatial correspondence of these clusters with canonical functional networks of the cerebral cortex was identified as the following: Right inferior frontal gyrus (frontoparietal), left precentral/middle frontal gyrus (dorsal attention), right precentral gyrus (somatomotor), precuneus (dorsal attention), left superior temporal gyrus (somatomotor), left/right lateral occipital cortex (visual), lingual gyrus (visual). Like the results of item memory enhancement, seed-based regression analysis again showed the involvement of widespread regions characteristic of distinct functional networks (**Figure 3**, lower panels). For instance, associative memory enhancement was negatively associated with functional connectivity between the sensory regions (occipital areas) and the dorsal attention network, along with functional connectivity between the more anterior association regions (left precentral/middle frontal gyrus, right inferior frontal gyrus) and the default mode network.

**Figure 3.**
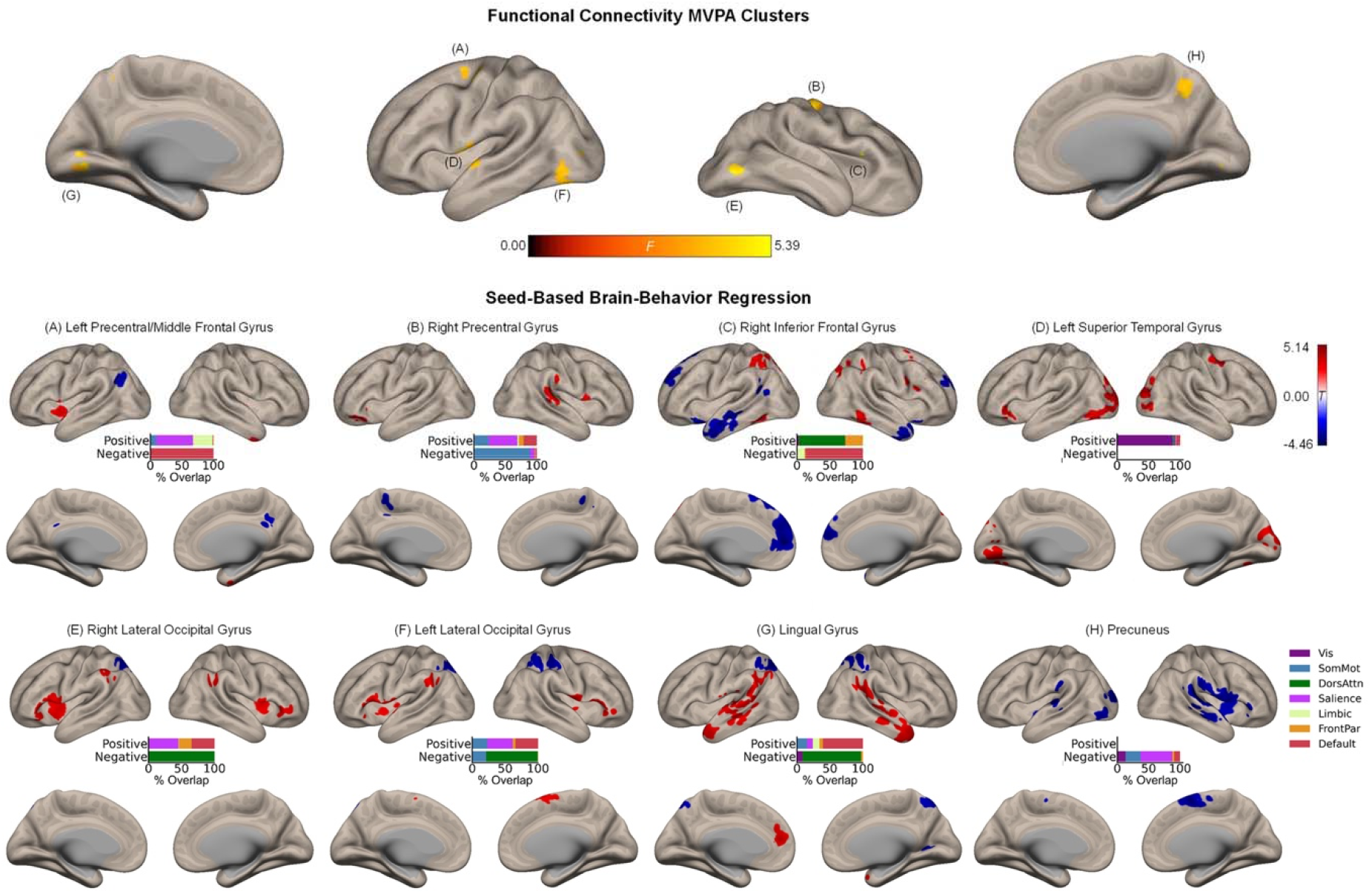
Intrinsic functional connectivity correlates of associative memory enhancement. Upper panel: Clusters identified in the functional connectivity MVPA as being significantly associated with associative memory enhancement. Lower panels: Whole-brain correlation maps between seed connectivity and associative memory enhancement. Bar graphs show the percentage of overlap between the areas identified in the correlation maps and canonical functional networks of the cerebral cortex (Yeo et al., 2011). Note that the voxel-wise intensity threshold for the correlation maps corresponds to |*t*| = 3.12 (*p* ≤ .001).

Beyond the cerebral cortex, associative memory enhancement was related to functional connectivity patterns of a few subcortical structures. Specifically, associative memory enhancement was positively associated with functional connectivity observed between the following seed-target pairs: The left precentral/middle frontal gyrus and a cluster in the left pallidum extending into the thalamus as well as between the same seed region and a cluster in the right posterior insula extending into the right putamen; the right precentral gyrus and a cluster in the right putamen as well as between the same seed region and a cluster spanning the posterior portion of the cerebellar cortex including Crus I, Crus II, and lobules VII- IX; and the left superior temporal gyrus and bilateral clusters in lobule VI of the cerebellar cortex (**Supplementary Figure S3**).

### Low dimensional decomposition of brain-behavior correlations

To facilitate the interpretation of our brain-behavior regression results, we performed low dimensional decomposition of spatial similarity observed across 11 second-level brain-behavior regression maps. Specifically, we first computed a matrix of η^2^ based on these inputs, which was submitted to PCA. This analysis revealed that the first two components together explained more than 80% of variance in the data (**Figure 4A**). To further characterize these two components in relation to the MVPA seed clusters, we visualized the PC scores of each region for Component 1 versus Component 2 in a two-dimensional scatter plot (**Figure 4B**). Furthermore, for each component, we calculated the sum of all contributing brain-behavior regression maps weighted by the respective PC scores (**Figure 4C**). These maps revealed the dominant axes along which the topography of brain-behavior relationships can be summarized.

**Figure 4.**
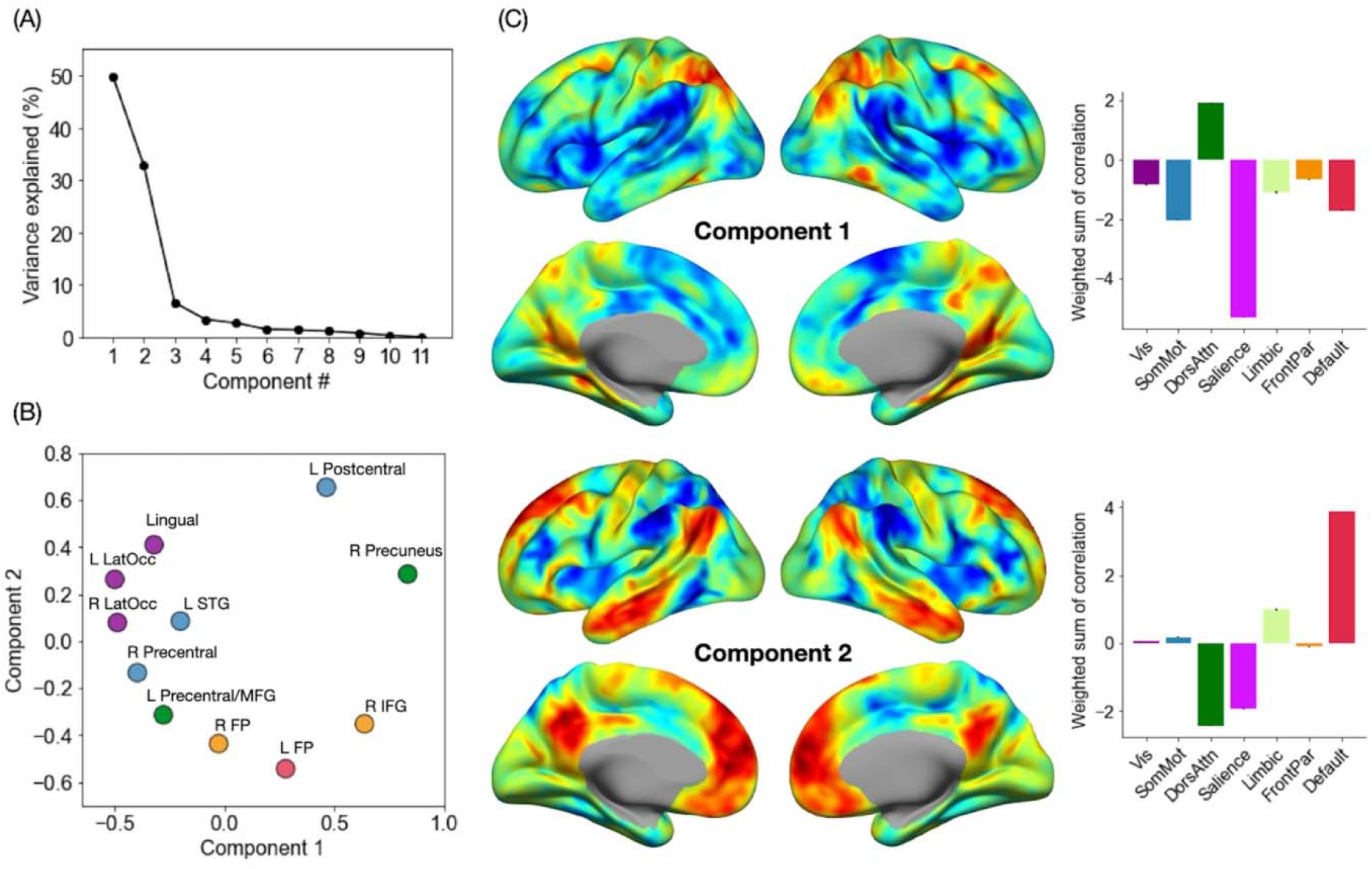
Low dimensional decomposition of brain-behavior correlations reveals two dominant axes of organization. (A) The percentage of variance in the data explained by each component. Note a clear knee in the curve showing variance explained by each component, suggesting that the first two components capture the most relevant aspects of brain-behavior correlation patterns. (B) PC scores of each MVPA seed cluster for the first two components. Each cluster is colored by the corresponding functional network label as determined by maximal spatial overlap with canonical network boundaries (Yeo et al., 2011). (C) Sums of all 11 correlation maps weighted by the respective PC scores. Bar graphs on the right show the breakdown of these voxel-wise weighted sum scores by the seven canonical networks of the cerebral cortex.

Component 1 primarily distinguished between regions that are part of the dorsal attention network and those that are part of the salience network. Specifically, the affective enhancement of memory was associated with functional connectivity of the dorsal attention network (e.g., superior parietal cortex, posterior middle temporal gyrus), positively with the prefrontal regions (left frontal pole, right inferior frontal gyrus) and negatively with the occipital regions (bilateral lateral occipital areas, lingual gyrus). Likewise, the affective enhancement of memory was also associated with functional connectivity of the salience network (e.g., insula, supramarginal gyrus), positively with sensorimotor regions scoring low on Component 1 and negatively with other parietal regions (precuneus, left postcentral gyrus). Further, Component 2 adds another dimension to this organization of brain-behavior relationships by distinguishing between regions that are part of the default mode network and those that are part of the dorsal attention and salience networks (the spectrum captured by Component 1). Specifically, the affective enhancement of memory was associated with functional connectivity of regions that are part of the default mode network, positively with sensorimotor regions (left postcentral, left lateral occipital cortex, lingual gyrus) and negatively with the prefrontal association regions. Likewise, the affective enhancement of memory was associated with functional connectivity of the dorsal attention/salience networks, positively with the prefrontal association regions and negatively with sensorimotor regions. In either component, clusters identified from analyses of item and associative memory enhancement were not clearly separated. Collectively, these findings point to the role of intrinsic connectivity of heteromodal association (i.e., dorsal attention, salience, and default mode) networks in supporting the affective enhancement of episodic memory.

## Discussion

The affective enhancement of episodic memory has been investigated extensively using task-related brain imaging. While these studies are useful in characterizing the associated neural mechanisms, the relationship between this phenomenon and intrinsic functional properties of the brain remained unclear. In this study, we utilized a whole-brain data-driven approach to the analysis of functional connectivity and found that the affective enhancement of memory was associated with intrinsic connectivity patterns of spatially distributed brain regions belonging to several functional networks in the cerebral cortex. These connectivity patterns in turn were primarily characterized by the involvement of heteromodal association (dorsal attention, salience, and default mode) networks as well as select subcortical structures. These findings suggest that the affective enhancement of episodic memory may be characterized as a whole-brain phenomenon, possibly supported by intrinsic functional interactions across several networks and structures in the brain.

To our knowledge, this is the first study linking the affective enhancement of episodic memory performance and intrinsic functional connectivity estimated from the same participants. However, our results identifying brain-behavior relationships in an ensemble of widespread regions are not surprising when considered in light of available neuroscientific evidence on varieties of arousal-related phenomena. Whether or not measures of memory are involved, affective arousal and the associated neural mechanisms are most typically studied in humans by presenting them with evocative stimuli and comparing brain activity patterns associated with these highly arousing versus neutral stimuli. Prior meta-analyses of such task-related neuroimaging studies revealed that processing of affective arousal involves a distributed network of cortical and subcortical structures, with frequently activated regions including portions of the default mode network (e.g., medial prefrontal cortex, ventrolateral prefrontal cortex) and the salience network (e.g., anterior mid-cingulate cortex, anterior insula, amygdala, hypothalamus) (Lindquist et al., 2016; Satpute et al., 2015). Our results are consistent with these findings, showing that intrinsic connectivity of these (cortical) regions was related to the degree to which episodic memory was enhanced by affective arousal across participants. Further, our results are also in line with prior work demonstrating that trait affective tendencies (reflecting mixtures of arousal and valence) were associated with intrinsic functional connectivity of widespread cortical areas including regions that are part of the default mode and salience networks as well as sensorimotor areas (Rohr et al., 2013).

Beyond the cerebral cortex (isocortex), we additionally demonstrated the relationship between the affective enhancement of memory and patterns of functional connectivity involving clusters identified in the striatum (i.e., putamen and pallidum), thalamus, and cerebellum. The involvement of these regions is consistent with their role in processing of affective arousal in general (Ceravolo et al., 2021; Lindquist et al., 2016; Satpute et al., 2019) as well as the enhancement of episodic memory by affective arousal more specifically (Dahlgren et al., 2020). Contributions of the cerebellum are particularly noteworthy given converging evidence identifying cerebellar involvement in broad aspects of affective processing (Adamaszek et al., 2017), going beyond its established role in sensorimotor domains. Because affective arousal is related to interoceptive sensations (Barrett et al., 2004), which emerge from changes in heart rate, glucose levels, inflammation, and so on, the current results are also consistent with our recent proposal that the cerebellum may play a domain-general computational role in support of the cerebral cortex, primarily concerned with efficient (i.e., predictive) regulation of the body’s internal systems (Katsumi et al., 2022; Katsumi, Kamona, et al., 2021). Collectively, these findings underscore the notion that large-scale intrinsic functional interactions across the whole brain may form the basis for the affective enhancement of episodic memory.

One notable observation is that the involvement of allocortical and subcortical structures within the medial temporal lobe was not observed in the current study. This is in contrast to the findings of task-related studies that consistently emphasized the role of such structures (e.g., the amygdala and the hippocampus) in the affective enhancement of episodic memory (Dahlgren et al., 2020; Murty et al., 2010). There are a few methodological factors that have possibly contributed to these seemingly discrepant results. For instance, our voxel-wise analyses were performed using a statistical threshold corrected for multiple comparisons at the cluster level within the whole brain search space. However, task-related studies examining smaller structures like the medial temporal areas commonly perform statistical analyses within a restricted search space, using relaxed thresholds, and/or in conjunction with various forms of small volume correction. While we do not rule out the possibility that intrinsic functional connectivity of the medial temporal lobe structures is related to the affective enhancement of memory, more targeted analytical approaches may be useful in identifying such (possibly more subtle) effects.

Additionally, our sample was based on a population-derived cohort from across the adult lifespan (Shafto et al., 2014), with more than 70% of our participants being older than 40 years of age. Aging is associated with reduced task-related functional connectivity changes in the medial temporal lobe linked to the affective enhancement of memory encoding (St. Jacques et al., 2009). Therefore, it is possible that greater representation of older adults in the current sample may have contributed to the attenuation of effects in the medial temporal lobe.

Across analyses, our results did not reveal distinct patterns of results between item and associative memory enhancement. This suggests that, at the level of intrinsic functional connectivity examined at the current threshold, the affective enhancement of these two types of memory is associated with partially overlapping networks of regions. However, it is important to note that the effect of affective arousal on associative memory appears to be sensitive to experimental designs, and that the degree of enhancement (or even impairment) is dependent on factors related to stimulus features, task instructions, and consolidation processes (e.g., retention interval) (reviewed in Dolcos, Katsumi, Denkova, et al., 2017). In the current study, both item and associative memory accuracy were enhanced by affective arousal, likely due to the instructions to “unitize” an item and the associated context in each trial (Chiu et al., 2013). Future work should examine similar brain-behavior associations using data acquired from experimental designs that are known to yield opposing effects of affective arousal on item and associative memory (e.g., Bisby et al., 2016). This would allow investigators to compare patterns of intrinsic functional connectivity associated with diverging effects of affective arousal on two types of memory.

A few limitations of this study should be noted. First, because we here focused on the effect of affective arousal in general, it remains unclear how valence may interact with the pattern of results observed in this study. Future work should expand on the present findings by comparing the degree of memory enhancement by positive and negative stimuli that are matched for arousal. Second, the sample collected for rating the image stimuli was relatively small. Investigators may benefit from the ratings that are replicated and expanded upon using larger samples in the future. Third, the cross-sectional nature of this work leaves unclear how reliable the observed effects are over time within individuals. This is an especially important area of future research in light of recent evidence suggesting that test-retest reliability of the affective enhancement of memory was low – i.e., confidence intervals of test-retest correlations across participants falling consistently below *r* = .5 (Schümann et al., 2020). Given that these authors used intentional encoding of stimuli, future studies involving incidental encoding should assess the replicability of both behavioral and neural effects associated with the affective enhancement of memory. Finally, given the spatial resolution of fMRI data, we were unable to examine smaller subcortical and brainstem structures (e.g., hypothalamus, periaqueductal gray, parabrachial nucleus), which are key to processing of affective arousal (Satpute et al., 2019). Future work should employ high-field functional imaging to help clarify the contributions of these regions.

Notwithstanding these limitations, the findings of this study—emerging from a highly demographically diverse, population-representative cohort—advance our understanding of the relationship between the affective enhancement of episodic memory and the intrinsic functional organization of the brain. Although it awaits independent replication and assessments of generalizability, the new evidence identified herein may be a first step toward possible ways to enhance affective well-being through modulation of intrinsic functional connectivity patterns that are associated with enhanced memory for subjectively arousing information.

## Authors Contributions

YK conceptualized the research; YK and MM performed data analysis and interpreted the results; YK wrote the first draft of the manuscript; MM wrote sections of the manuscript; All authors contributed to manuscript revision and approved the submitted version.

## Acknowledgement

During the preparation of this manuscript, MM was supported by the Department of Veterans Affairs, Office of Academic Affiliations, California War Related Illness and Injury Study Center (WRIISC-CA) fellowship program. The authors thank the Cambridge Centre for Ageing and Neuroscience consortium for their generous open sharing of the dataset.

