## Supplementary Materials and Methods for "Affective enhancement of episodic memory is associated with widespread patterns of intrinsic functional connectivity across the adult lifespan"

### Katsumi & Moore: Supplementary Materials and Methods

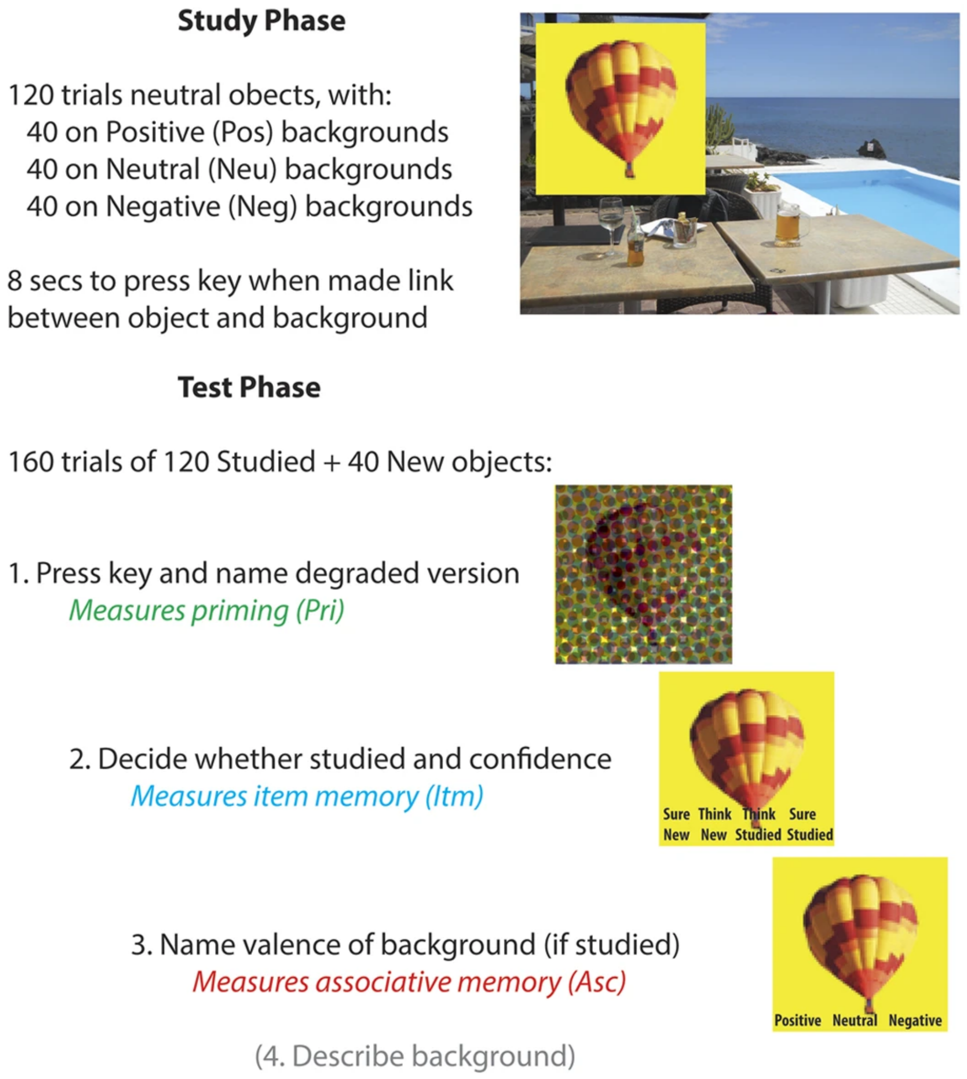


**Supplementary Figure S1.** Summary of Study and Test phases. Figure reproduced from Henson et al. (2016), with permission.

*Justification of the number of components in functional connectivity MVPA*

Functional connectivity MVPA requires the user to specify the number of principal components ($N_c$) to use in characterizing between-participants variability in whole-brain voxel-to-voxel connectivity patterns. Convention in the literature is to define $N_c$ based on the participant-to-component ratio (Arnold Anteraper et al., 2019, 2020; Guell et al., 2020; Katsumi et al., 2021; Kawagoe et al., 2019; Takamiya et al., 2021; Whitfield-Gabrieli et al., 2016). In the current study, we initially performed first-level MVPA using 15 components, approximately corresponding to a 20:1 ratio between the number of participants and that of components. In addition to the participant-to-component ratio, recent work has begun taking into account the individual and cumulative explained variance for each component (Morris et al., 2021). It is important to keep in mind that the inclusion of too many components might (1) make it difficult to detect an effect of a given size across the components in the omnibus $F$-test, and (2) begin to explain variance in white matter voxel connectivity that is considered noise in this work.

**Supplementary Table S1** below shows the percentage explained variance in gray matter functional connectivity variability by each component as well as the cumulative percentage for each additional component. Beyond seven components, each additional component explains less than 1% of variance, with the total variance explained by the first seven components being ~92%. Based on these results, we reasoned that seven components would sufficiently represent the observed between-participants variability in whole-brain connectivity structure at each voxel.

| Component | Variance explained (%) | Cumulative variance explained (%) |
|---|---|---|
| 1 | 74.07 | 74.07 |
| 2 | 7.28 | 81.34 |
| 3 | 4.16 | 85.51 |
| 4 | 2.58 | 88.09 |
| 5 | 1.82 | 89.91 |
| 6 | 1.37 | 91.28 |
| **7** | **1.04** | **92.32** |
| 8 | 0.81 | 93.13 |
| 9 | 0.65 | 93.78 |
| 10 | 0.52 | 94.30 |
| 11 | 0.43 | 94.74 |
| 12 | 0.36 | 95.10 |
| 13 | 0.30 | 95.40 |
| 14 | 0.25 | 95.65 |
| 15 | 0.22 | 95.87 |

**Supplementary Table S1.** Individual and cumulative variance in gray matter connectivity variability explained by each MVPA component.

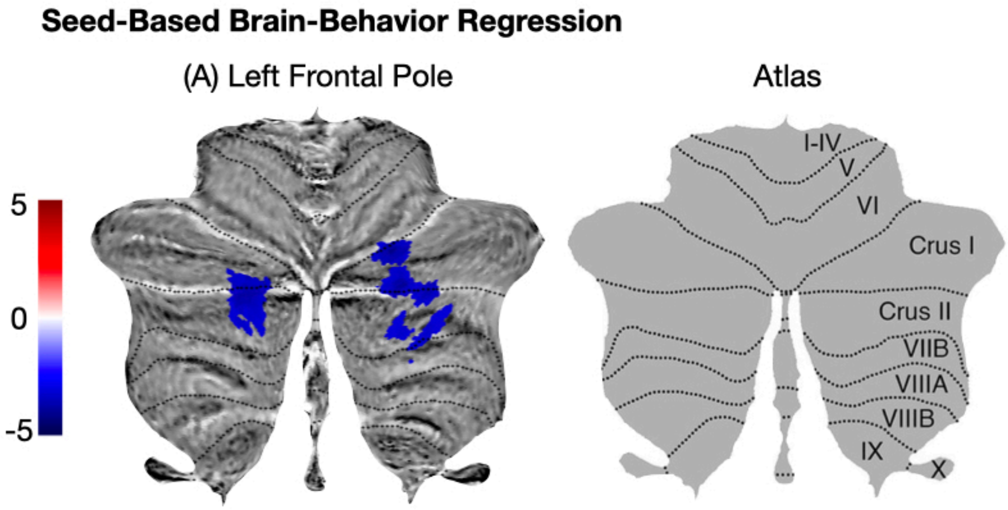


**Supplementary Figure S2.** Intrinsic functional connectivity correlates of item memory enhancement in the subcortex. Left panel: Clusters of suprathreshold voxels in the cerebellar cortex whose functional connectivity with the left frontal pole seed (see **Figure 2** in the main text) was negatively associated with item memory enhancement. These clusters are depicted on a flat map representation of the cerebellar cortex (Diedrichsen & Zotow, 2015). Right panel: Subregions of the cerebellar cortex depicted on a flat map. Reproduced from Guell et al. (2018), with permission.

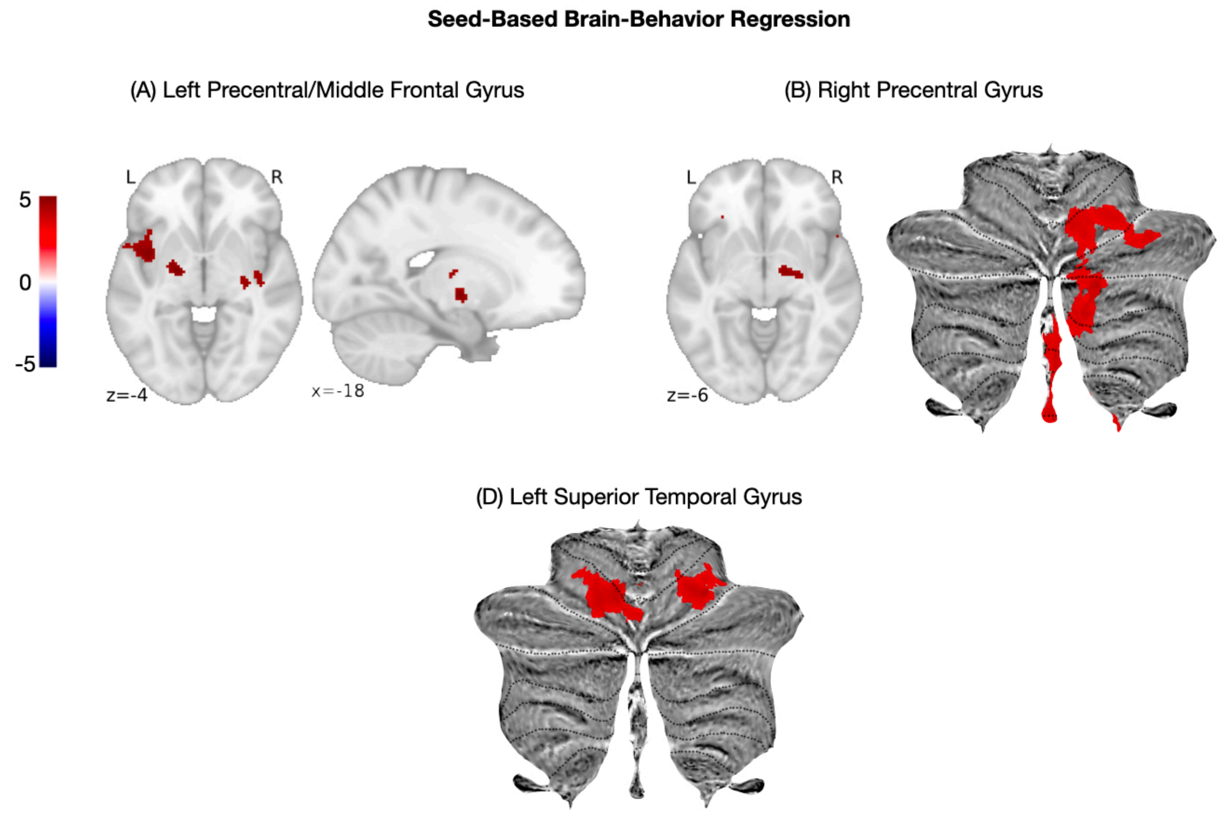


**Supplementary Figure S3.** Intrinsic functional connectivity correlates of associative memory enhancement in the subcortex. These volumetric and cerebellar flat map projections identify clusters of suprathreshold voxels whose functional connectivity with the seed regions (see **Figure 3** in the main text) was positively associated with associative memory enhancement.
